# A fast and efficient smoothing approach to Lasso regression and an application in statistical genetics: polygenic risk scores for chronic obstructive pulmonary disease (COPD)

**DOI:** 10.1101/2020.03.06.980953

**Authors:** Georg Hahn, Sharon M. Lutz, Nilanjana Laha, Michael H. Cho, Edwin K. Silverman, Christoph Lange

## Abstract

High dimensional linear regression problems are often fitted using Lasso approaches. Although the Lasso objective function is convex, it is not differentiable everywhere, making the use of gradient descent methods for minimization not straightforward. To avoid this technical issue, we apply Nesterov smoothing to the original (unsmoothed) Lasso objective function. We introduce a closed-form smoothed Lasso which preserves the convexity of the Lasso function, is uniformly close to the unsmoothed Lasso, and allows us to obtain closed-form derivatives everywhere for efficient and fast minimization via gradient descent. Our simulation studies are focused on polygenic risk scores using genetic data from a genome-wide association study (GWAS) for chronic obstructive pulmonary disease (COPD). We compare accuracy and run-time of our approach to the current gold standard in the literature, the FISTA algorithm. Our results suggest that the proposed methodology provides estimates with equal or higher accuracy than the FISTA algorithm while having the same asymptotic runtime scaling. The proposed methodology is implemented in the R-package *smoothedLasso*, available on the Comprehensive R Archive Network (CRAN).

## 1 Introduction

Many substantive research questions in health, economic and social science require solving a classical linear regression problem *y* = *Xβ* + *ϵ*. Here, the data matrix *X* ∈ ℝ^*n×p*^, the parameter vector *β* ∈ ℝ^*p*^, and the response vector *y* ∈ ℝ^*n*^ encode *n* ∈ ℕ linear equations in *p* ∈ ℕ variables, and *ϵ* ∼ *N*_*n*_(0, *σ*^2^*I*_*n*_) is an *n*-dimensional, independently and normally distributed error term with mean zero and some variance *σ*^2^ *>* 0 (where *I*_*n*_ denotes the *n*-dimensional identity matrix). The approach remains one of the most widely used statistical analysis tools.

Traditionally, linear regression problems have been solved by finding parameter estimates *β* that minimize the squared error, leading to the least squares estimate arg 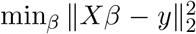. However, their lack of robustness, as well as sparsity requirements in high dimensional settings with *p* ≫ *n*, are two problems that have led to the development of alternative estimation approaches, e.g. the *least absolute shrinkage and selection operator* (Lasso) of Tibshirani (1996) or *least-angle regression* (LARS) of Efron et al. (2004).

This article focuses on Lasso regression, which obtains regression estimates 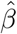 by solving

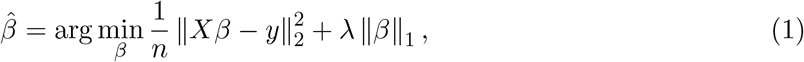

where ‖ · ‖ _2_ is the Euclidean norm, ‖ · ‖ _1_ is the *L*_1_ norm, and *λ >* 0 is a tuning parameter (called the Lasso regularization parameter) controlling the sparseness of the solution 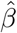.

As the objective function in eq. (1) is convex, minimization via steepest descent (quasi-Newton) methods is sensible. However, many applications in biostatistics, especially those that are focused on *big data*, such as the simultaneous analysis of genome-wide association studies (Wu et al., 2009) or the calculation of polygenic risk scores (Mak et al., 2016), involve data sets with several thousand parameters and are often sparse. In such applications, conventional gradient-free solvers can lose accuracy. This is due to the non-differentiability of the *L*_1_ penalty term in eq. (1), i.e. the term 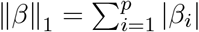.

We address this issue by smoothing the Lasso objective function. We apply Nesterov smoothing (Nesterov, 2005) to the non-differentiable ‖ *β* ‖_1_ penalty term in eq. (1). This will result in an approximation of the Lasso objective function that is differentiable everywhere. The Nesterov formalism depends on a smoothing parameter *µ* that controls the smoothness level.

Our approach has three major advantages: (1) The smoothing preserves the convexity of the Lasso objective function; (2) it allows us to obtain closed-form derivatives of the smoothed function everywhere which we use in a gradient descent algorithm; and (3) it provides uniform error bounds on the difference between the smoothed and the original objective functions. The error bounds depend only on the smoothing parameter *µ*, and by letting *µ* → 0, the smoothed Lasso objective function can be made arbitrarily close to the original Lasso objective function, thus justifying its use as a surrogate of the original Lasso objective. However, when smoothing the *L*_1_ penalty of the Lasso, the resulting regression estimates are typically dense. That is, the lack of the *L*_1_ penalty somewhat causes a loss of the variable selection property, though regression estimates for those variables which are non-selected by the classical Lasso will typically be very small. Therefore, an easy fix to obtain sparse estimates again is to apply some form of thresholding.

We benchmark our proposed smoothing approach against the current gold standard for minimizing the Lasso objective function, the FISTA algorithm of Beck and Teboulle (2009). FISTA is a proximal gradient version of the algorithm of Nesterov (1983) which combines the basic iterations of the Iterative Shrinkage-Thresholding Algorithm (Daubechies et al., 2004) with a Nesterov acceleration step. Among others, the algorithm is implemented in the R-package *fasta* on CRAN (Chi et al., 2018). We use this package as a benchmark. The FISTA algorithm requires the separate specification of the smooth and non-smooth parts of the objective function including their explicit gradients. In contrast to our approach, a variety of tuning parameters need to be selected by the user, e.g. an initial starting value, an initial stepsize, parameters determining the *lookback* window for non-monotone line search, and a shrinkage parameter for the stepsize.

The benchmark of our approach against FISTA is carried out on simulated data, with the aim to assess accuracy and runtime scaling of both algorithms in *n* and *p*, and on a real-world dataset of polygenic risk scores of COPDGene (genetic epidemiology of COPD). Polygenic risk scores are a statistical aggregate of risks typically associated with a set of established DNA variants. The goal of polygenic risk scores is to predict the disease risk of an individual, that is the susceptibility to a certain disease given the individual’s genotypes at the established risk loci and other important epidemiological covariates such as age, sex etc. Such scores are usually calibrated on large genome-wide association studies (GWAS) via high-dimensional regression of the genetic variants to the outcome. Since the potential for broad-scale clinical use to identify people at high risk for certain diseases has been demonstrated (Khera et al., 2018), polygenic risk scores have become a widespread tool for the early identification of patients who are at high risk for a certain disease and who could benefit from intervention measures. For this reason, increasing the accuracy of scores is desirable, which is what we address with the proposed smoothing approach.

This article is structured as follows. Section 2 applies Nesterov smoothing to the Lasso objective function. Using polygenic risk scores as an example, we evaluate the proposed methodology in a simulation study in Section 3. The article concludes with a discussion in Section 4. The proof of our theoretical result is given in Appendix C. The methodology of this article is implemented in the R-package *smoothedLasso*, available on CRAN (Hahn et al., 2020).

Throughout the article, *X*_*·,i*_ denotes the *i*th column of a matrix *X*. Similarly, *X*_*I,·*_ (and *y*_*I*_) denote the submatrix (subvector) consisting of all rows of *X* (entries of *y*) indexed in the set *I*. Moreover, *X*_*−I,·*_ (and *y*_*−I*_) denote the submatrix (subvector) consisting of all rows of *X* (entries of *y*) not indexed in the set *I*. Finally, | · | denotes the absolute value, and ‖ · ‖ _*∞*_ denotes the supremum norm.

## 2 Smoothing the Lasso objective function

This section lays the theoretical foundation for the modified Lasso approach we propose to address the non-differentiability of the *L*_1_ penality term in eq. (1), while the smooth *L*_2_ term remains unchanged. The substitute of the term 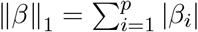 in eq. (1) will be computed with the help of a technique called Nesterov smoothing, whose details are given in Section 2.1. In Section 2.2, we simplify the general Nesterov smoothing approach for the particular case of the Lasso penalty and show how our approach results in explicit closed-form expressions for both the smoothed Lasso and its gradient. The uniform closeness of the smoothed Lasso we propose and the original Lasso objective function is proven in Section 2.3.

### 2.1 Summary of Nesterov smoothing

We briefly summarize the idea of Nesterov smoothing. We are given a piecewise affine and convex function *f* : ℝ^*q*^ → ℝ which we aim to smooth, where *q* ∈ ℕ. We assume that *f* is composed of *k* ∈ ℕ linear pieces (components) and can thus be expressed as

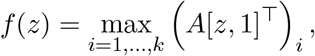

where *A* ∈ ℝ^*k×*(*q*+1)^ is a matrix whose rows contain the linear coefficients for each of the *k* linear pieces (with the constant coefficients being in column *q* + 1), *z* ∈ ℝ^*q*^, and [*z*, 1] ∈ ℝ^*q*+1^ denotes the vector obtained by concatenating *z* and the scalar 1.

Let *µ* ≥ 0 be the Nesterov smoothing parameter. Using a so-called proximity (or prox) function *ρ*, which is assumed to be nonnegative, continuously differentiable, and strongly convex, *f* is replaced by the approximation

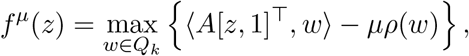

where *Q*_*k*_ ⊆ ℝ^*k*^ is the unit simplex in *k* dimensions defined as

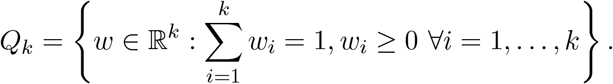

As shown in (Nesterov, 2005, Theorem 1), *f*^*µ*^ is both smooth and uniformly close to *f*, meaning that for all *z* ∈ ℝ^*q*^, the approximation error is uniformly upper bounded by

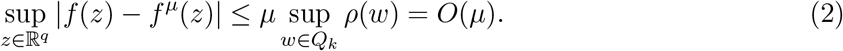

The larger the value of *µ* the more *f* is smoothed, while *µ* = 0 recovers the original function *f* = *f* ^0^. Two choices of the prox function are given a closer look in Section 2.2. Smoothing with the so-called entropy prox function results in the smooth approximation 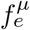 having a closed-form expression given by

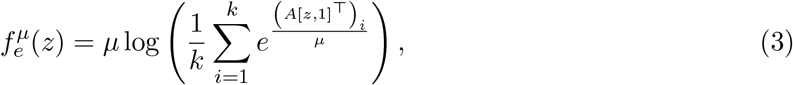

which satisfies the uniform bound

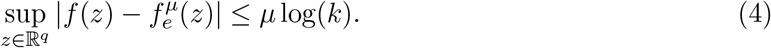

Similarly, the smooth approximation of *f* with the help of the so-called squared error prox function given by 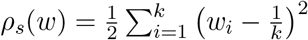 for *w* ∈ ℝ^*k*^ can be written as

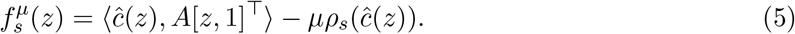

In eq. (5), *ĉ* (*z*) ∈ ℝ^*k*^ is the Michelot projection (Michelot, 1986) of the vector *c*(*z*) = (*c*_1_(*z*), *…, c*_*k*_(*z*)), given componentwise by *c*_*i*_(*z*) = 1*/µ* · (*A*[*z*, 1]^⊤^)_*i*_ − 1*/k* for *i* ∈ {1, *…, k*}, onto the *k*-dimensional unit simplex *Q*_*k*_. The approximation 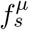 via squared error prox function satisfies the uniform bound

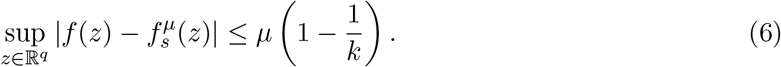

### 2.2 Application to the Lasso objective function

For given *X* ∈ ℝ^*n×p*^, *y* ∈ ℝ^*n*^, and *λ >* 0, according to eq. (1), the Lasso objective function *L* : ℝ^*p*^ → ℝ given by

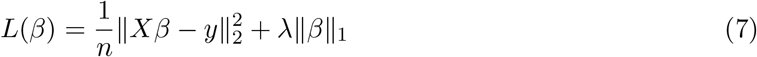

is smooth in its first term but non-differentiable in the penalty term 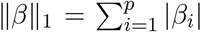. We thus smooth the latter, where it suffices to apply Nesterov smoothing to each absolute value independently.

Let *k* = 2. Using one specific choice of the matrix *A* ∈ ℝ^2*×*2^ given by

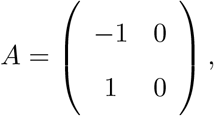

we rewrite the (one dimensional) absolute value as *f* (*z*) = max{−*z, z*} = max_*i*=1,2_ (*A*[*z*, 1]^⊤^) _*i*_, where here and in the following subsections *z* ∈ ℝ is a scalar.

#### 2.2.1 Entropy prox function

For the entropy prox function, eq. (3) with *A* as in Section 2.2 simplifies to

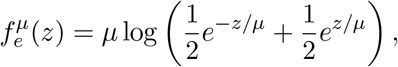

which according to eq. (4) satisfies the approximation bound

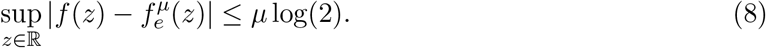

The first and second derivatives of 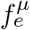 are given by

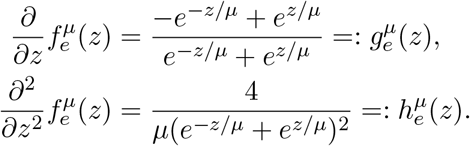

Together, smoothing eq. (7) with the entropy prox function results in

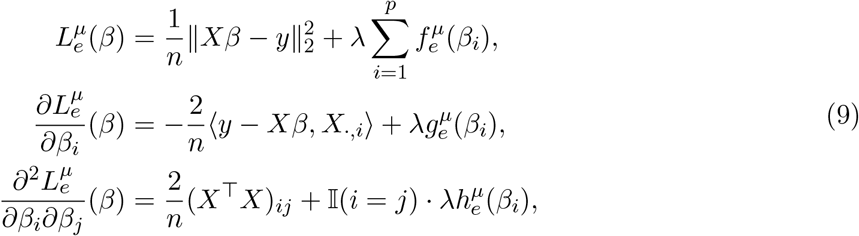

where the entropy prox smoothed Lasso is 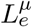, its explicit gradient is 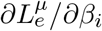, and its Hessian matrix is given by 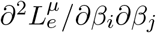. The function 𝕀 (·) denotes the indicator function.

In principle, the Lasso objective can be minimized using a (second order) Newton-Raphson or a (first order) quasi-Newton approach. However, since *X* ∈ ℝ^*n×p*^ with *n < p*, the matrix *X*^⊤^*X* is singular, meaning that for the Hessian to be invertible one needs the added diagonal elements 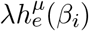 to be large. However, this is usually not true, since if *β*_*i*_ is nonzero, then in a neighborhood of the true Lasso estimate the term 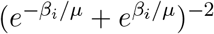 will be exponentially small. Thus to make 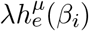 large for a fixed *µ*, we need *λ* to be exponentially large. Likewise, given *λ* and *β*_*i*_, too small or too large values of *µ* will make 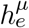 vanish. However, since typically *λ* and *µ* are fixed, the Hessian in eq. (9) will be singular except for a few artificial cases and thus the second order Newton-Raphson method will not be applicable. In the simulations we therefore focus on quasi-Newton methods which require only 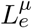 and its gradient 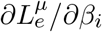.

#### 2.2.2 Squared error prox function

Similarly, eq. (5) with *A* as in Section 2.2 simplifies to

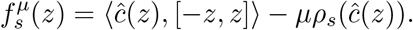

where *ĉ* (*z*) ∈ ℝ^2^ is the Michelot projection of the vector *c*(*z*) = 1*/µ* · [−*z, z*] − 1*/k* onto the two-dimensional unit simplex *Q*_2_. According to eq. (6), we obtain the approximation bound

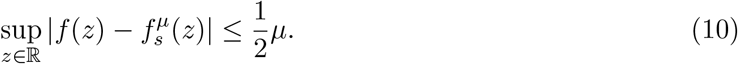

The derivative of 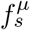 is given by

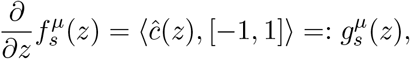

see (Hahn et al., 2017, Lemma 4) for a proof of this result. The second derivative 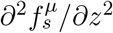 does not have a closed form expression, though it can be approximated numerically. Analogously to the results for the entropy prox function, smoothing eq. (7) with the squared error prox function results in

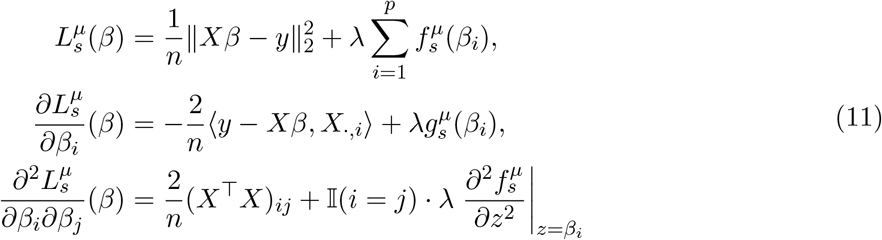

where as before, the smoothed Lasso via squared error prox function is 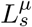, its explicit gradient is 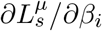, and its Hessian matrix is 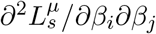.

As in Section 2.2.1 we observe that the Hessian matrix is singular since *X*^⊤^*X* is singular, and since the additional diagonal entries stemming from 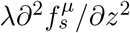 are usually too small to make the Hessian invertible. We therefore again resort to a quasi-Newton method to minimize 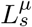 with the help of its gradient 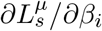 only.

### 2.3 Theoretical error bounds

The approximation bounds on the absolute value, given in eqs. (8) and (10), carry over to bounds on the distance between the unsmoothed and smoothed Lasso objective functions, as shown in the following proposition.

#### Proposition 1

*The smoothed Lasso objective functions with entropy and squared error prox, given in* eqs. (9) *and* (11), *satisfy the following uniform bounds:*

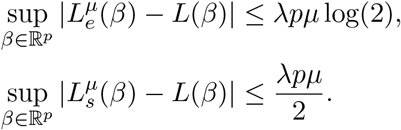

*Moreover, both* 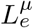 *and* 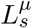 *are strictly convex*.

In Proposition 1 the Lasso parameter *λ >* 0 and the dimension *p* are fixed for a particular estimation problem, thus allowing to make the approximation error arbitrarily small as the smoothing parameter *µ* → 0. This justifies the use of our proposed smoothed Lasso as a surrogate of the original (non-smooth) Lasso objective function. Moreover, the convexity of the Lasso is preserved.

## 3 Simulation studies for polygenic risk scores for COPD

In this section, we evaluate three approaches to compute Lasso estimates on simulated data (Section 3.1) as well as real data coming from a genome-wide association study for COPDGENE (Regan et al., 2010), in which polygenic risk scores are computed and evaluated (Section 3.2). Of the three approaches we consider, the first two utilize existing methodology, while the last approach implements the methodology we developed in this paper:

1. We carry out the minimization of eq. (1) using R’s function *optim*. The *optim* function implements the quasi-Newton method *BFGS* for which we supply the explicit (though non-smooth) gradient *∂L/∂β*_*i*_ = −2*/n* ·⟨ *y* − *Xβ, X*_*·,i*_ ⟩ + λsign(*β*_*i*_). This approach will be referred to as the *unsmoothed Lasso*;
2. We use the FISTA algorithm as implemented in the *fasta* R-package (Chi et al., 2018), available on *The Comprehensive R Archive Network* (R Core Team, 2014);
3. We minimize the smoothed Lasso objective function of eq. (9) using its explicit gradient. Note that due to the smoothed *L*_1_ penalty, the resulting estimates returned by the smoothed Lasso are typically dense, though sparse estimates are easily obtained by applying some form of thresholding.

The main function of the *fasta* package which implements the FISTA algorithm, also called *fasta*, requires the separate specification of the smooth and non-smooth parts of the objective function including their explicit gradients. We follow Example 2 in the *vignette* of the *fasta* R-package in Chi et al. (2018) and supply both as specified in eq. (7). Additionally, we employ a uniform random starting value as done for the unsmoothed and smoothed Lasso. The initial stepsize is set to *τ* = 10 as in Example 2 of Chi et al. (2018). The lookback window for non-monotone line search and the shrinkage parameter for the stepsize are left at their default values.

Three potential implementations are unconsidered in this simulation section for the following reasons. The *glmnet* algorithm of Friedman et al. (2010), available in the R-package *glmnet*, is a variant of FISTA which performs a cyclic update of all coordinates, whereas FISTA updates all coordinates per iteration. We thus focus on FISTA. Since the R-package *SIS* accompanying Fan and Li (2001) does itself rely on *glmnet* for computing regression estimates, we omit it in this section. The LARS algorithm of Efron et al. (2004) is implemented in the R-package *lars* on CRAN (Hastie and Efron, 2013). As remarked in Friedman et al. (2010), LARS is slower than glmnet/ FISTA. Additionally, since the implementation of Hastie and Efron (2013) always computes a full Lasso path, it is considerably slower than the other methods.

All results are averages over 100 repetitions. The Lasso regularization parameter was fixed at *λ* = 0.3 after performing 10-fold cross validation. Details of the cross validation computation are given in Appendix A. The smoothing parameter of the approach of Section 2.2 was fixed at *µ* = 0.1 in all simulations. As shown in Proposition 1, the smaller *µ* the better the proposed smoothed Lasso will approximate the original Lasso, though numerical instabilities may occur if *µ* is chosen too close to the machine precision. All computations were carried out on 100 cluster nodes, each one having a lx24-amd64 architecture with 4 cores per node, 94 GB of memory, and running 64-bit CentOS 6 Linux.

### 3.1 Application to simulated data

We designed our simulation study so that it mimics the application to a genome-wide association study, i.e. we simulate *X* ∈ ℝ^*n×p*^ from a multidimensional normal distribution, where the entries of the mean vector of the multidimensional normal distribution are sampled independently from a uniform distribution in [0, 0.5]. To ensure positive definiteness of the covariance matrix Σ of the multidimensional normal distribution, we set 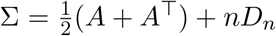, where *D*_*n*_ is a *n* × *n* matrix with ones on its diagonal, and zeros otherwise. The added term *nD*_*n*_ ensures positive definiteness.

After *X* is obtained, we generate the entries of the true *β* independently from a standard normal distribution, and set all but *nz* ∈ {0, *…, p*} out of the *p* entries to zero. The number of non-zero entries *nz* ∈ ℕ is a parameter of the simulations. The response *y* ∈ ℝ^*n*^ is then easily obtained as *y* = *Xβ* + *ϵ*, where the entries of the noise vector *ϵ* ∈ ℝ^*n*^ are generated independently from a Normal distribution with mean zero and some standard deviation *σ*. The smaller the standard deviation, the easier the recovery of *β* will be (see Appendix B for a study on how the choice of *σ* influences the recovery of *β*). We will employ *σ* = 0.5 in all simulations. In this subsection, we fix the number of true non-zero parameters at 20% (that is, *nz* = 0.2*p*), resulting in sparse parameter vectors.

Figure 1 (left) shows results on simulated data of dimension *n* ∈ [1, 10000] while keeping *p* = 1000 fixed. We measure the accuracy of the obtained Lasso estimates through their *L*_2_ norm to the generated true parameters. We observe that the unsmoothed and smoothed Lasso approaches seem to suffer from numerical instabilities for small *n*. As *n* increases, both the unsmoothed and smoothed Lasso approaches stabilize. The FISTA algorithm yields stable estimates for all *n*. The accuracy of the smoothed Lasso approaches the one of FISTA for large *n*. As expected, all methods become more accurate as the number of data points *n* increases.

**Figure 1.**
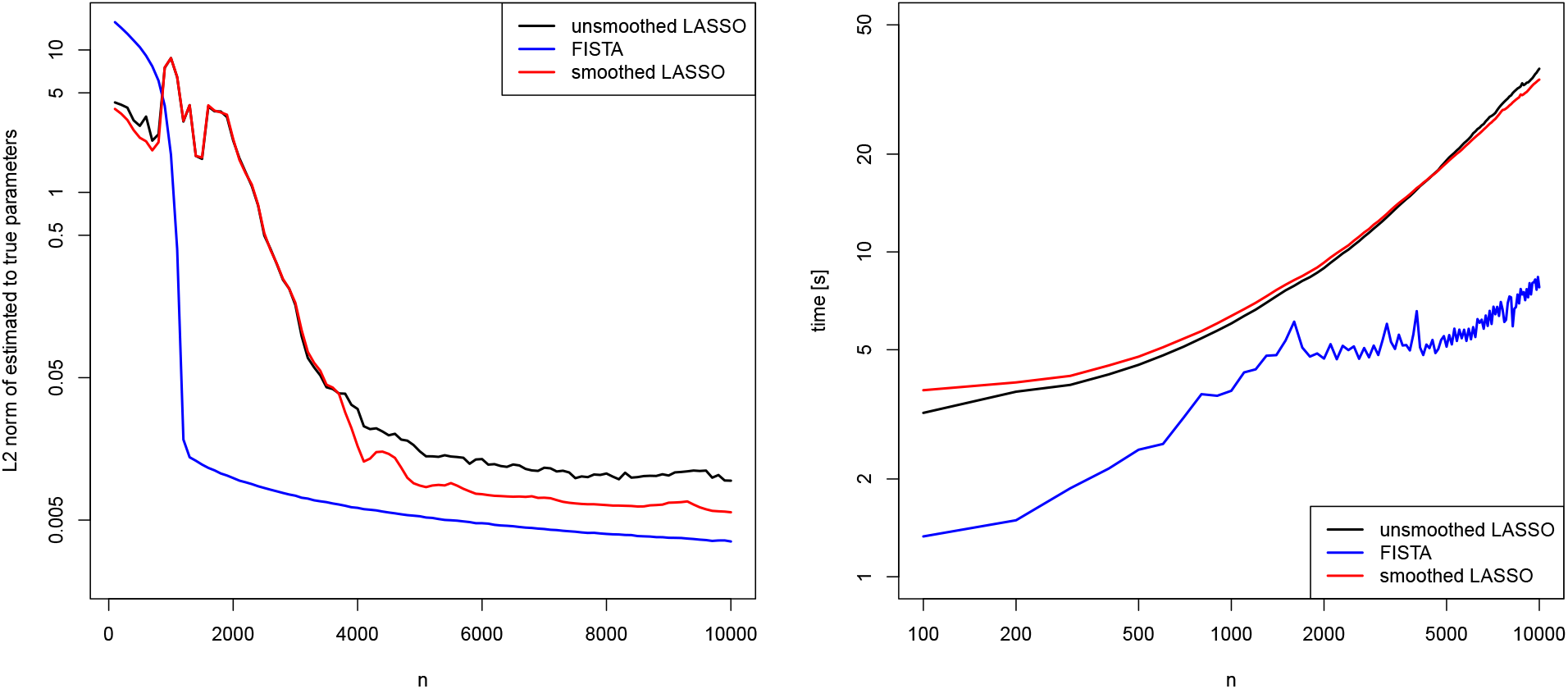
: Simulation study: *L*_2_ norm of parameter estimate to truth (left) and runtime in seconds (right) as a function of *n* ∈ [1, 10000] while *p* = 1000. Logarithmic scale on the y-axes.

Figure 1 (right) shows that the unsmoothed and smoothed Lasso approaches have an almost identical runtime. We find that although highly optimized, FISTA is only a low multiple factor faster than our approaches and, more importantly, its asymptotic runtime scaling is comparable to the one of the smoothed Lasso.

Similarly to the previous experiment, in Figure 2 (left) we keep *n* = 1000 fixed and investigate the dependence of all four approaches on *p* ∈ [1, 5000]. Again, the unsmoothed and smoothed Lasso approaches seem to suffer from numerical instabilities for smaller *p*, while FISTA is very stable. Importantly, the smoothed Lasso seems to outperform FISTA for very large *p*. As expected, while keeping the data size *n* fixed, estimation becomes more challenging for all methods as *p* increases.

**Figure 2.**
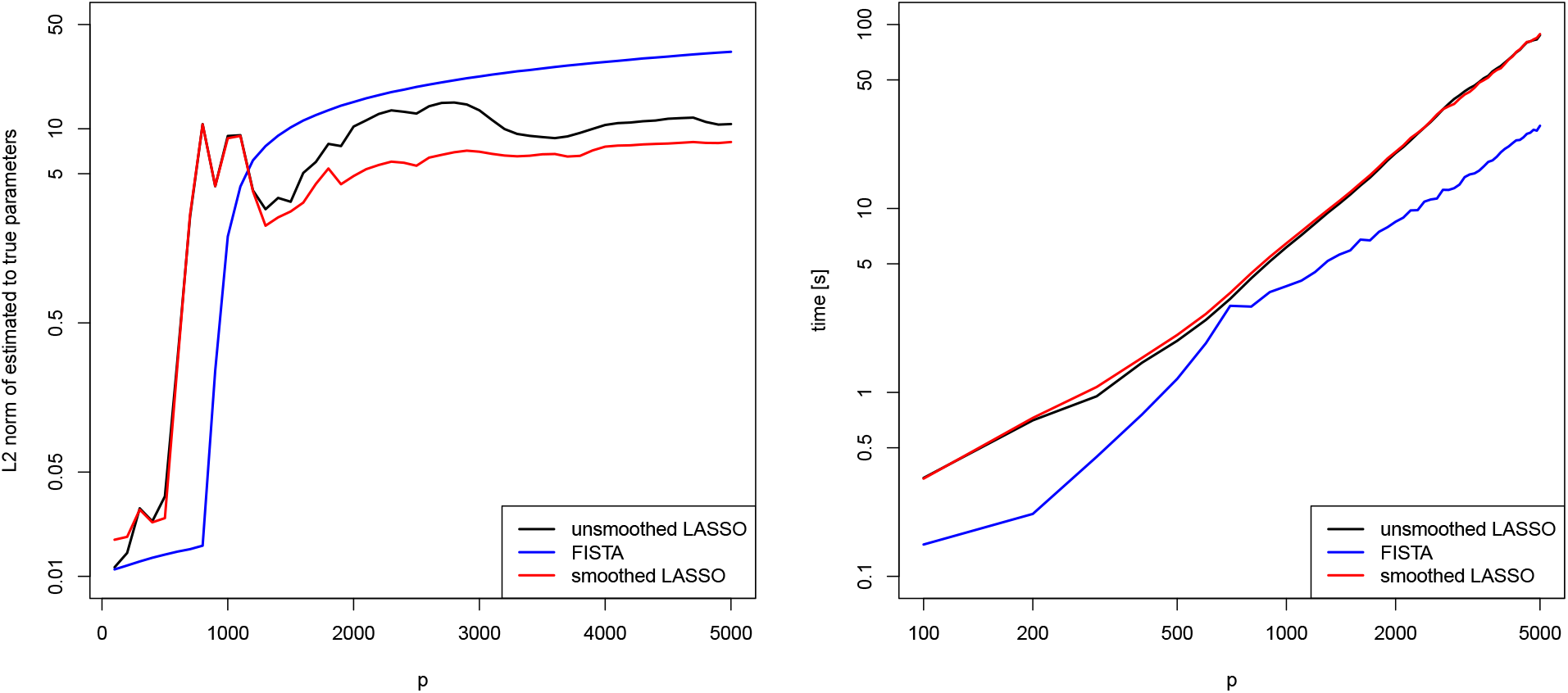
: Simulation study: *L*_2_ norm of parameter estimate to truth (left) and runtime in seconds (right) as a function of *p* ∈ [1, 5000] while *n* = 1000. Logarithmic scale on the y-axes.

Figure 2 (right) confirms the timing results seen in the assessment of the dependence on *n*. The unsmoothed and smoothed approaches have virtually equal speeds, while FISTA is again slightly faster. Notably, the scaling of the runtimes seems to be roughly equal for all three methods.

Since the unsmoothed Lasso does not come with the guarantee we established for our smoothing approach (see Section 2.3) and is outperformed by the smoothed Lasso, we will focus in the remainder of the simulations on FISTA and smoothed Lasso only.

### 3.2 Application to polygenic risk scores

We evaluate the smoothed Lasso approach on polygenic risk scores of the COPDGene study (genetic epidemiology of COPD), a multi-center case-control study designed to identify genetic determinants of COPD and COPD-related phenotypes (Regan et al., 2010). The study has been sequenced as part of the TOPMed Project. The data is available through NHLBI TOPMed (2018).

For the study, non-Hispanic Whites and African Americans aged 45 to 80 were recruited as COPD cases and controls. The dataset contains *n* = 4010 individuals (rows), all of which had at least 10 pack-years of smoking history. For each individual, we observe *p* = 9153 datapoints, among them the covariates *age, sex, packyears, height* and five PCA vectors. The remaining entries are SNP data per individual. The input data are summarized in a matrix *X* ∈ ℝ^*n×p*^. The response *y* ∈ ℝ^*n*^ is the *fev1* ratio, also called the Tiffeneau-Pinelli index, per individual. It describes the proportion of lung volume that a person can exhale within the first second of a forced expiration in a spirometry (pulmonary function) test. We aim to fit the four covariates, the five PCA vectors, and the SNP data to the response using our smoothing approach, with the aim to obtain a polygenic risk score of higher accuracy than obtained through conventional methods (see Section 1).

#### 3.2.1 Results from a single run

We solve *y* = *Xβ* for the given *X* and *y* using our smoothed Lasso approach of eq. (9), and compare it to the FISTA algorithm.

Table 1 shows results for a single application of both algorithms. After computing the estimate 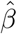 with each method, we consider 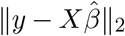, the *L*_2_ norm between the fitted and generated (true) response. We observe that FISTA with standard choices of its tuning parameters seems to have trouble locating the minimum of the Lasso objective function, and is thus worse than smoothed Lasso. However, the FISTA method beats our approach in terms of runtime.

**Table 1:** COPDGene study: *L*_2_ norm of fitted to true response and runtime in seconds for a single application of any method to the dataset of polygenic risk scores.

| method | $L_2$ norm | runtime [s] |
| --- | --- | --- |
| FISTA | 284.6 | 66.6 |
| smoothed Lasso | 26.3 | 116.6 |

#### 3.2.2 Cross validation

To quantify the accuracy of the methods further, we perform a simple cross validation experiment in which we withhold a random set of row indices *I* (of varying size) of the dataset *X* and the corresponding entries of the response *y* and fit a linear model to the rest of the data, that is we fit *y*_*−I*_ = *X*_*−I,·*_*β*. After obtaining an estimate 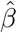, we use the withheld rows of *X* to predict the withheld entries of *y*, that is we compute 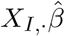. We evaluate the quality of the prediction by computing the *L*_2_ norm 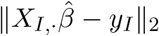 between predicted and withheld data.

Figure 3 (left) shows results of this cross-validation experiment. We observe that FISTA with default choices of its tuning parameters seems to have difficulties to converge to the minimum of the Lasso objective function, whereas smoothed Lasso performs better. Not surprisingly, the quality of the prediction becomes worse in general as the number of withheld entries increases, since predictions are based on fewer and fewer datapoints. However, this decrease in accuracy seems to be more pronounced for FISTA than for our smoothed Lasso.

**Figure 3.**
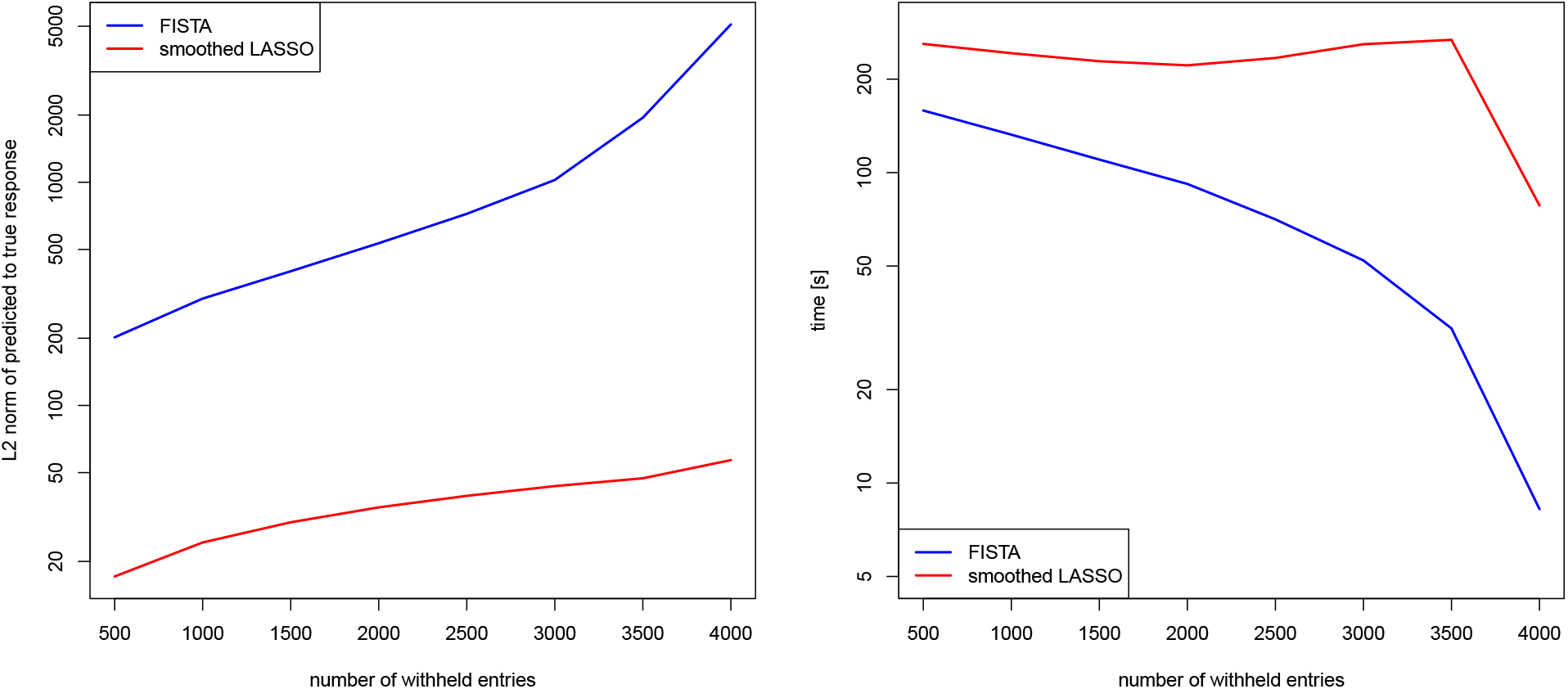
: COPDGene study: *L*_2_ norm of predicted to withheld data in cross-validation (left) and runtime in seconds (right) as a function of the number of withheld entries. Dataset of polygenic risk scores. Logarithmic scale on the *y*-axes.

Finally, Figure 3 (right) displays runtime measurements for both approaches. We observe that smoothed Lasso seems to be rather insensitive to the number of withheld entries apart from a very large number of withheld entries. The FISTA algorithm is the faster method, however it exhibits a greater sensitivity to the number of withheld entries, that is the size of the estimation problem.

### 3.3 Application to synthetic polygenic risk scores

We aim to extend the simulations of Section 3.2 in order to vary the sparsity of the parameter estimate 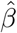. To this end, we change the simulation setting as follows. Leaving *X* ∈ ℝ^*n×p*^ unchanged (*n* = 4010 individuals and *p* = 9153 datapoints), we simulate a parameter vector *β* ∈ ℝ^*p*^ in which *nz* ∈ {0, *…, p*} entries are drawn from a Beta distribution with shape parameters 1.5 and 10. This will produce nonzero entries in the vector *β* of magnitude around 0.15, which is realistic in practice. The remaining *p* − *nz* entries are set to zero. We then calculate the response as *y* = *Xβ* + *ϵ*, where the entries of the noise vector *ϵ* ∈ ℝ^*n*^ are generated independently from a Normal distribution with mean 0 and standard deviation 0.1.

After generating *X* and *y*, we again use FISTA and smoothed Lasso to recover an estimate 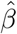. We evaluate the quality of the estimate using 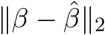, that is using the *L*_2_ norm between truth and estimate.

Figure 4 (left) shows results as a function of the number *nz* of non-zero entries in the generated true *β*. We observe that in this experiment, and in contrast to Figures 1 and 2, the smoothed Lasso yields both more accurate and more stable estimates than FISTA (as expressed through a lower deviation from the truth in *L*_2_ norm).

**Figure 4.**
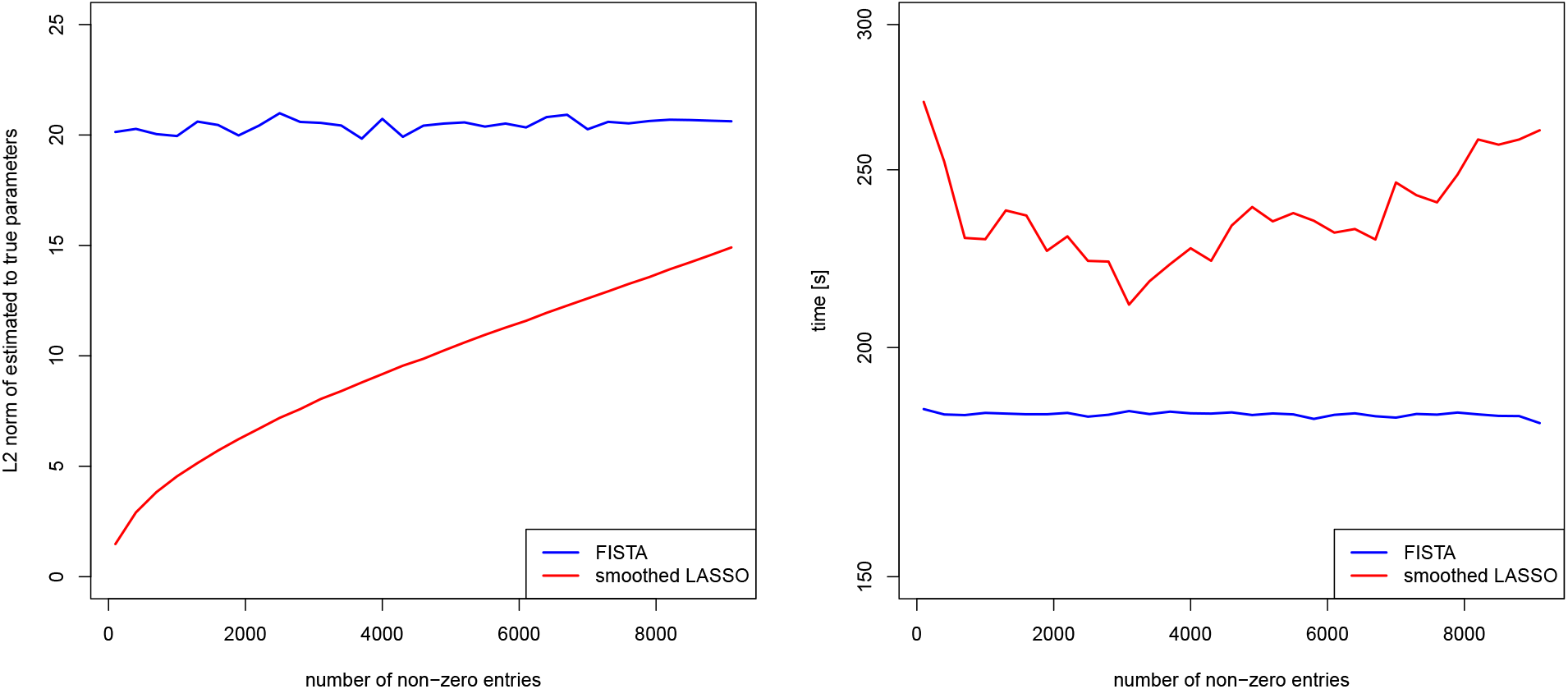
: COPDGene study: *L*_2_ norm of parameter estimate to truth (left) and runtime in seconds (right) as a function of the number of non-zero entries in the simulated true parameter vector. Dataset of polygenic risk scores. Logarithmic scale on the *y*-axis of the time plot.

Figure 4 (right) shows runtime results for both approaches, showing that in this experiment the runtime scaling of FISTA seems to be rather insensitive to the simulation scenario. Our smoothed Lasso is slower than FISTA, and its runtime is more volatile. We note that in this experiment, the almost constant runtime of FISTA, irrespective of the simulation scenario, is likely a result of its inability to produce stable estimates in the first place.

Overall, we conclude from the simulations that in the scenarios we investigated, our smoothed Lasso approach yields estimates with equal or improved accuracy in comparison to the gold standard FISTA, exhibits roughly the same runtime scaling as FISTA, and comes with a guarantee on its accuracy.

## 4 Discussion

This article investigated a smoothing approach for penalized regression using the Lasso. The smoothing approach allows us to obtain smooth gradients everywhere, which facilitate minimization of the convex but non-smooth Lasso objective function.

Importantly, our approach comes with a guarantee of correctness. We prove a uniform bound on the distance between the unsmoothed and smoothed Lasso objective function, thus justifying the use of our smoothed Lasso as a surrogate of the original (non-smooth) Lasso objective function. In the scenarios that we investigated, our simulation results suggest that the proposed smoothing approach yields Lasso estimates with equal or improved accuracy than the gold standard in the literature, the FISTA algorithm of Beck and Teboulle (2009). Our approach can directly be utilized for the computation of more accurate polygenic risk scores via high-dimensional regression of genetic variants from genome-wide association studies (and other covariates) to a certain outcome. This has the potential to increase the precision of risk prediction, thereby potentially identifying patients at high risk early on when they are still able to benefit from preventive steps targeted at reducing their chance of getting the disease.

Moreover, the idea of smoothing is not restricted to Lasso regression. Other important and widespread regression operators, such as the elastic net or graphical Lasso, might benefit from a similar smoothing approach to increase the accuracy of the obtained estimates. Extending the methodology to those operators remains for future work.

## A Selection of the Lasso regularization parameter via cross validation

We aim to select the Lasso regularization parameter *λ* using cross validation. To this end, for the simulation scenario described in Section 3.1 (in particular, for the chosen noise level of *σ* = 0.5 and the sparsity level of 20%, as well as *n* = 1000 and *p* = 100), we perform 10-fold cross validation as described in Tibshirani (2013).

To be precise, we first fix a grid of admissible values of *λ* from which we would like to choose the regularization parameter (here, *λ* ∈ {0, 0.05, 0.1, 0.15, *…*, , 1}). We then randomly divide the *n* data points into *K* = 10 disjoint sets (folds) *I*_1_, *…, I*_*K*_ such that 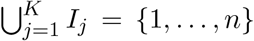. For each *j* ∈ {1, *…, K*}, we withhold the indices in *I*_*j*_ and fit a linear model 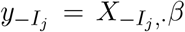 using the FISTA algorithm. After obtaining an estimate 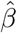, we use the withheld rows of *X* indexed in *I*_*j*_ to predict the withheld entries of *y*, that is we compute 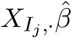. We evaluate the accuracy of the prediction with the *L*_2_ norm, that is we compute 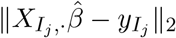. Repeating this computation for all *j* ∈ {1, *…, K*} allows us to compute an average *L*_2_ error over the *K* folds (called the *cross validation error*). We plot this error as a function of *λ*.

The result is shown in Figure 5. We observe that for the simulation scenario we consider in Section 3.1, the choice *λ* = 0.3 is sensible.

**Figure 5:**
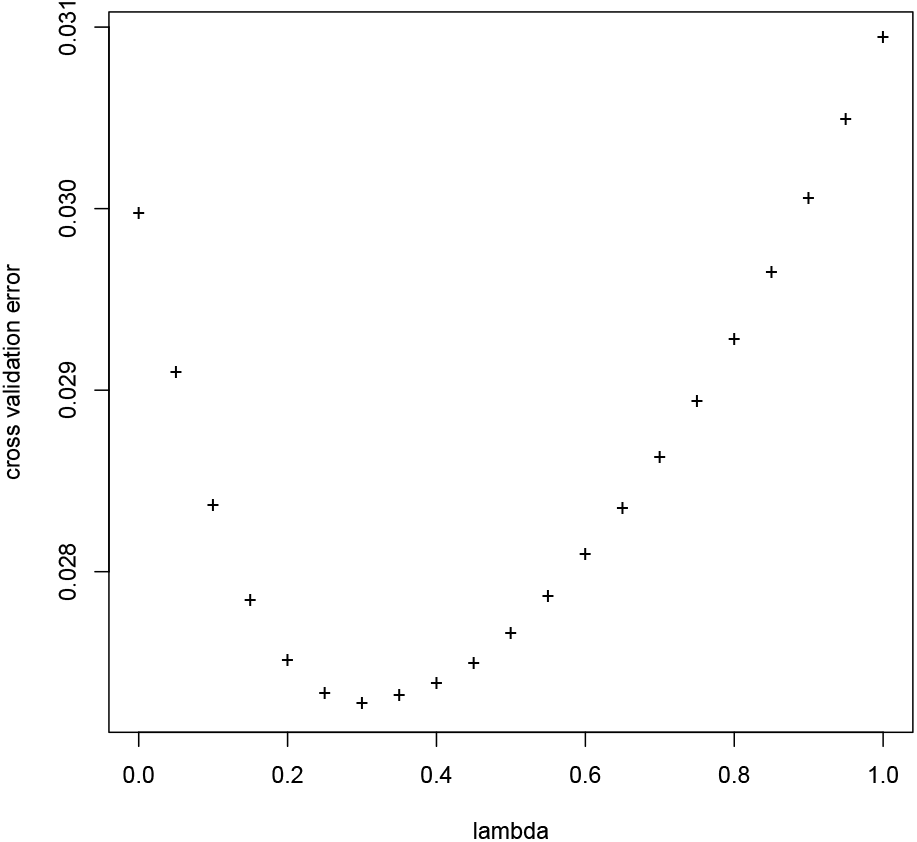
Cross validation error as a function of the Lasso regularization parameter *λ* for the FISTA algorithm. Simulated data with *n* = 1000 and *p* = 100.

## B Sensitivity analysis

In the linear regression model *y* = *Xβ* + *ϵ* under consideration in this work (see Section 1), it is easy to see that the larger the noise/error *ϵ*, the harder it will be to obtain accurate estimates of *β*.

To quantify this statement, Figure 6 presents a sensitivity analysis for the recovery accuracy of the parameter estimate *β* (measured as the *L*_2_ norm between the fitted parameter estimate returned by the unsmoothed Lasso, FISTA algorithm, and smoothed Lasso, to the truth) as a function of the standard deviation *σ*. The setup of the simulation is identical to the one of Section 3.1, though now *n* = 100 and *p* = 200 are fixed. The entries of the noise vector *ϵ* ∈ ℝ^*n*^ in the model *y* = *Xβ* + *ϵ* are generated independently from a Normal distribution with mean zero and a varying standard deviation *σ* ∈ [0, 10].

**Figure 6:**
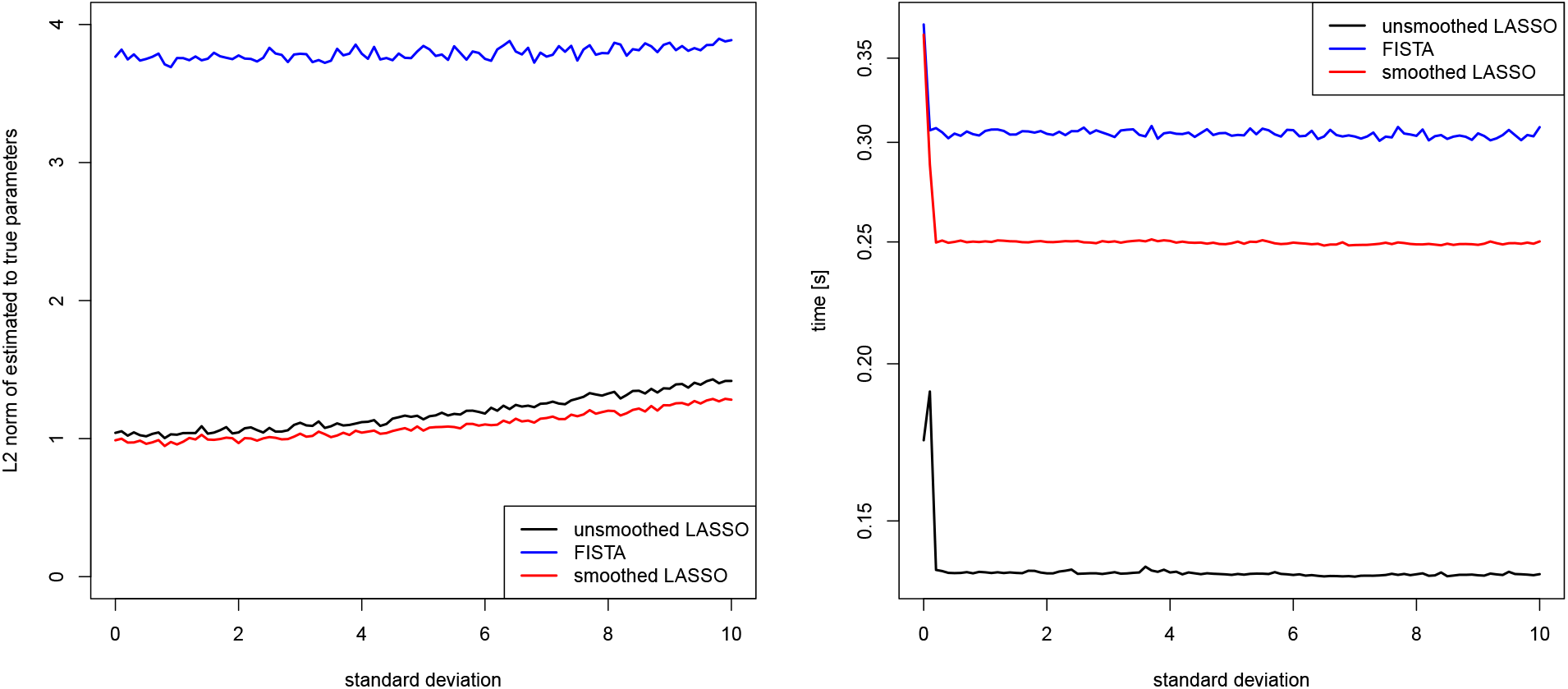
Sensitivity analysis on simulated data: *L*_2_ norm of parameter estimate to truth (left) and runtime in seconds (right) as a function of the standard deviation *σ* while *n* = 1000 and *p* = 2000. Logarithmic scale on the y-axis of the right plot.

Figure 6 (left) shows that, as expected, the accuracy of the recovered estimate of *β* decreases for all methods as *σ* increases. However, this increase seems rather slow. The runtime as a function of *σ*, depicted in Figure 6 (right), stays roughly constant for all methods, as expected.

## C Proof of Proposition 1

*Proof*. The bounds on 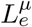 and 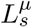 follow from eq. (8) and eq. (10) after a direct calculation. In particular, for the entropy prox function,

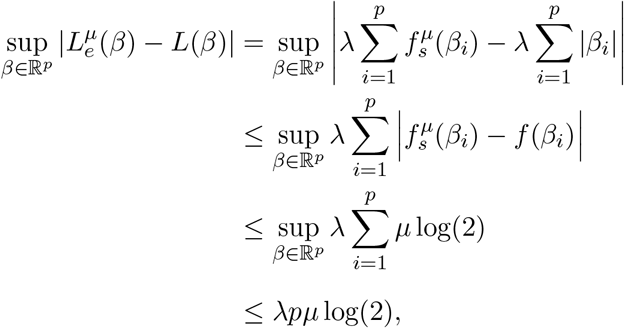

where *f* is as defined in Section 2.2 and where it was used that *λ* ≥ 0. The result for the squared error prox smoothed 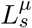 is proven analogously.

Since both 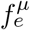 and 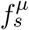 are convex according to (Nesterov, 2005, Theorem 1), and since the least squares term 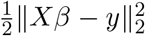 is convex, it follows that both 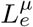 and 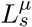 remain convex as the sum of two convex functions.

To be precise, strict convexity holds true. Observe that the second derivative of the entropy smoothed absolute value of Section 2.2.1 is given by

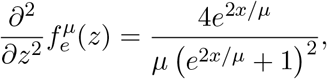

which is always positive, thus making 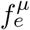 strictly convex. Therefore, 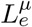 is strictly convex as the sum of a convex function and a strictly convex function. Similar arguments show that 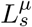 is strictly convex. □

## Notes

### Competing Interest Statement

The authors have declared no competing interest.

## References

Beck, A. and Teboulle, M. (2009). A Fast Iterative Shrinkage-Thresholding Algorithm for Linear Inverse Problems. SIAM J Imaging Sciences, 2(1):183–202.

Chi, E., Goldstein, T., Studer, C., and Baraniuk, R. (2018). fasta: Fast Adaptive Shrink-age/Thresholding Algorithm. R-package version 0.1.0.

Daubechies, I., Defrise, M., and Mol, C. (2004). An iterative thresholding algorithm for linear inverse problems with a sparsity constraint. Comm. Pure Appl. Math., 57(11):1413–1457.

Efron, B., Hastie, T., Johnstone, I., and Tibshirani, R. (2004). Least angle regression. Ann Stat, 32(2):407–499.

Fan, J. and Li, R. (2001). Variable Selection via Nonconcave Penalized Likelihood and its Oracle Properties. J Am Stat Assoc, 96(456):1348–1360.

Friedman, J., Hastie, T., and Tibshirani, R. (2010). Regularization Paths for Generalized Linear Models via Coordinate Descent. Journal of Statistical Software, 33(1):1–22.

Hahn, G., Banerjee, M., and Sen, B. (2017). Parameter Estimation and Inference in a Continuous Piecewise Linear Regression Model.

Hahn, G., Lutz, S. M., Laha, N., and Lange, C. (2020). smoothedLasso: Smoothed LASSO Regression via Nesterov Smoothing. R-package version 1.3: https://cran.r-project.org/package=smoothedLasso.

Hastie, T. and Efron, B. (2013). lars: Least Angle Regression, Lasso and Forward Stagewise. R-package version 1.2.

Khera, A. V., Chaffin, M., Aragam, K. G., Haas, M. E., Roselli, C., Choi, S. H., Natarajan, P., Lander, E. S., Lubitz, S. A., Ellinor, P. T., and Kathiresan, S. (2018). Genome-wide polygenic scores for common diseases identify individuals with risk equivalent to monogenic mutations. Nature Genetics, 50:1219–1224.

Mak, T., Porsch, R., Choi, S., Zhou, X., and Sham, P. (2016). Polygenic scores via penalized regression on summary statistics. Genet Epidemiol, 41(6):469–480.

Michelot, C. (1986). A finite algorithm for finding the projection of a point onto the canonical simplex of R^n^J Optimiz Theory App, 50(1):195–200.

Nesterov, Y. (1983). A method of solving a convex programming problem with convergence rate O(1/k2Dokl Akad Nauk SSSR, 269(3):543–547.

Nesterov, Y. (2005). Smooth minimization of non-smooth functions. Math. Program. Ser. A, 103:127–152.

NHLBI TOPMed (2018). Boston Early-Onset COPD Study in the National Heart, Lung, and Blood Institute (NHLBI) Trans-Omics for Precision Medicine (TOPMed) Program.

R Core Team (2014). R: A Language and Environment for Statistical Computing. R Foundation for Stat Comp, Vienna, Austria.

Regan, E., Hokanson, J., Murphy, J., Make, B., Lynch, D., Beaty, T., Curran-Everett, D., Silverman, E., and Crapo, J. (2010). Genetic epidemiology of copd (copdgene) study design 2. COPD, 7:32–43.

Tibshirani, R. (1996). Regression Shrinkage and Selection Via the Lasso. J Roy Stat Soc B Met, 58(1):267–288.

Tibshirani, R. (2013). Model selection and validation 1: Cross-validation.

Wu, T., Chen, Y., Hastie, T., Sobel, E., and Lange, K. (2009). Genome-wide association analysis by lasso penalized logistic regression. Bioinformatics, 25(6):714–721.

